# In vivo CRISPR screening identifies geranylgeranyl diphosphate as a pancreatic cancer tumor growth dependency

**DOI:** 10.1101/2024.05.03.592368

**Authors:** Casie S. Kubota, Stephanie L. Myers, Toni T. Seppälä, Richard A. Burkhart, Peter J. Espenshade

## Abstract

Cancer cells must maintain lipid supplies for their proliferation and do so by upregulating lipogenic gene programs. The sterol regulatory element-binding proteins (SREBPs) act as modulators of lipid homeostasis by acting as transcriptional activators of genes required for fatty acid and cholesterol synthesis and uptake. SREBPs have been recognized as chemotherapeutic targets in multiple cancers, however it is not well understood which SREBP target genes are essential for tumorigenesis. Using parallel in vitro and in vivo CRISPR knockout screens, we identified terpenoid backbone biosynthesis genes as essential for pancreatic ductal adenocarcinoma (PDAC) tumor development. Specifically, we identified the non-sterol isoprenoid product of the mevalonate pathway, geranylgeranyl diphosphate (GGPP), as an essential lipid for tumor growth. Mechanistically, we observed that restricting mevalonate pathway activity using statins and SREBP inhibitors synergistically induced apoptosis and caused disruptions in small G protein prenylation that have pleiotropic effects on cellular signaling pathways. Finally, we demonstrated that *geranylgeranyl diphosphate synthase 1* (*GGPS1*) knockdown significantly reduces tumor burden in an orthotopic xenograft mouse model. These findings indicate that PDAC tumors selectively require GGPP over other lipids such as cholesterol and fatty acids and that this is a targetable vulnerability of pancreatic cancer cells.

## Introduction

Lipids support cell growth and proliferation by serving as structural components of membranes, energy sources, signaling molecules, and protein modifications. Under conditions of nutrient stress, tissues adapt to maintain normal cellular function. Cellular lipid homeostasis is maintained by the sterol regulatory element-binding protein (SREBP) class of transcription factors (1,2). When cellular lipids are limiting, ER-resident SREBPs are trafficked to the Golgi by the SREBP cleavage activating protein (SCAP). SREBPs are then activated by proteolytic cleavage and upregulate the transcription of lipid metabolic genes responsible for cholesterol and fatty acid synthesis and uptake such as *low-density lipoprotein receptor (LDLR)*, *stearoyl-CoA desaturase (SCD)*, and *HMG-CoA reductase (HMGCR).* Two genes, *SREBF1* and *SREBF2*, encode three isoforms, SREBP-1a, SREBP-1c, and SREBP-2, with distinct gene programs. SREBP-1 is generally responsible for upregulating fatty acid synthesis, while SREBP-2 controls cholesterol synthesis and uptake (3). Lipid metabolic reprogramming has been implicated as an important facet of tumor metabolism, prompting the hypothesis that SREBPs may be an effective target for cancer therapy (4,5).

Pancreatic ductal adenocarcinoma (PDAC) is one of the deadliest forms of cancer with a 5-year survival rate of 12% (6). Poor patient prognosis is due to both late diagnosis as well as limited treatment options (7). *KRAS* oncogene driver mutations are present in >90% of pancreatic cancer cases, which is recognized as the initiating genetic event for PDAC development (8). Inactivating mutations in the tumor suppressor *TP53* are also found in >60% of pancreatic cancers and are known to promote PDAC disease progression (8). Functional consequences of these mutations have been characterized extensively and are known to be drivers of metabolic reprogramming within tumors. Namely, constitutively active *KRAS* is known to promote de novo lipid synthesis, and loss of function of *TP53* was shown to stimulate isoprenoid and cholesterol synthesis through upregulation of the mevalonate pathway (9–12). Together, these studies provide motivation for targeting lipogenesis through the SREBP pathway in PDAC.

Activation of the SREBP pathway has been shown to promote cancer progression (13–16). Recently, we demonstrated that *SCAP* is required for PDAC tumor growth in orthotopic and subcutaneous xenografts as well as in a genetically engineered PDAC mouse model (17). Given that *SCAP* is required for tumor growth and functions through SREBPs to upregulate a broad program of lipogenic genes, we proposed that SREBP target genes contribute to tumor growth. To identify SCAP-dependent genes required for tumor establishment, we designed a focused CRISPR knockout library against SREBP target genes and performed parallel in vitro and orthotopic in vivo CRISPR screens using a patient-derived *KRAS^G12D^/TP53^L344P^* PDAC cell line. Strikingly, both sets of genetic screens identified non-sterol isoprenoid synthesis as a PDAC cell vulnerability. Mechanistically, we observed that inhibiting flux through this synthesis pathway is detrimental for protein prenylation with pleiotropic effects on cellular signaling and protein localization, ultimately leading to apoptotic cell death. Taken together, these observations further support the pursuit of SREBPs as a therapeutic target for PDAC and illustrate the essentiality of the mevalonate pathway for PDAC tumor formation.

## Materials and Methods

### Cell Culture

Human pancreatic ductal adenocarcinoma cell lines Pa02c and Pa03c were derived from liver metastatic tumors, and Pa16c and Pa20c cells from primary tumors. Cell lines were generously provided by Dr. Anirban Maitra (Department of Pathology, Johns Hopkins University) (18). Genotyping verified the presence of *KRAS* and *TP53* mutations. Cells were maintained at 37°C and 5% CO_2_ in 2D monolayer culture and confirmed as *Mycoplasma* negative (MycoAlert Mycoplasma Detection Kit, Lonza #LT07-701). Cells were maintained in DMEM medium (Corning #10-013-CV) containing 4.5 g/L glucose, L-glutamine, and sodium pyruvate supplemented with 10% (v/v) fetal bovine serum (FBS, Gibco #10438026) and 1,000 U/mL penicillin-streptomycin (Gibco #15140122). Cells were lipid-starved using DMEM medium (Corning #10-013-CV) containing 4.5 g/L glucose, L-glutamine, and sodium pyruvate supplemented with 10% lipoprotein-deficient serum (LPDS, Kalen #880100-2) and 1,000 U/mL penicillin-streptomycin (Gibco #15140122) unless otherwise specified.

### Transfection and Lentivirus Transduction

HEK293T cells were co-transfected with psPAX2 (Addgene #12260), VSV.G (Addgene #14888), and transfer plasmids in a 45:5:50 ratio using PolyFect transfection reagent (Qiagen #301107) following the manufacturer’s protocol. Viral supernatant was passed through a 0.45 µm PVDF membrane filter and cells were transduced for 48 h with 10 µg/mL polybrene (Sigma #TR-1003-G).

### CRISPR Screening

Stable Cas9-expressing Pa03c cells were transduced with lentiCas9-Blast (Addgene #52962) lentivirus. Cells were selected with blasticidin and dilution cloned in 96-well plates to isolate monoclonal populations. Cas9 expression was evaluated by western blot. Cas9 cells were infected with library lentivirus pool at an MOI of ∼0.3. Library-infected cells were then split into 10% FBS or 10% LPDS culture medium for in vitro screen or injected into the pancreas of nude mice at 5 x 10^5^ cells in 50 µL 1:1 Matrigel:PBS (Corning #CB-40230, Corning #21-040-CV). For the in vitro screen, cells were passaged every 3 days for 12 days. Genomic DNA was collected using the DNeasy Blood and Tissue Kit (Qiagen #69504). For the in vivo screen, tumors were grown for 3 weeks and treated as separate samples. In vitro and in vivo PCR amplicons were sequenced together with initial library-infected cell samples as reference.

### Mouse Xenograft Studies

All animal experiments were conducted following protocols approved by the Johns Hopkins Institutional Animal Care and Use Committee (IACUC) and Johns Hopkins Research Animal Resources (RAR). 8-12 week old female athymic nude mice (Hsd:Athymic Nude-*Foxn1^nu^*, Envigo) were used for both subcutaneous and orthotopic xenografts. For the subcutaneous model, mice were anesthetized with isoflurane using a vaporizer and 0.5 x 10^6^ Pa03c or 1 x 10^6^ Pa16c cells in 0.1 mL 1:1 Matrigel:PBS (Corning #CB-40230, Corning #21-040-CV) were injected subcutaneously on the right flank. Length and width of tumors were measured using a digital caliper (Fowler) every 2-3 days in addition to mouse weight. Tumor volume was calculated according to the following: (minimum width^2^ x maximum length)/2 (19). Mice were administered Fluvastatin intraperitoneally (dissolved in water, 20 mg/kg) or by oral gavage (dissolved in water, 50 mg/kg) daily. For the orthotopic model, mice were anesthetized with isoflurane using a vaporizer, and a small incision was made into the left flank of the skin and abdominal wall. The pancreas was externalized, and 0.5 x 10^5^ cells in 20 µL 1:1 Matrigel:PBS (Corning #CB-40230, Corning #21-040-CV) were injected into the tail of the pancreas using a Hamilton syringe with a 30G needle. The abdominal wall and skin were sutured closed using 4-0 Monocryl Violet suture (Ethicon #Y464G). Buprenorphine-SR (Zoopharm #1Z-73000-190711) was administered at 1 mg/kg subcutaneously following surgery. Following euthanasia, tumors and/or whole pancreata were flash-frozen in liquid nitrogen and stored at -80°C or fixed in 10% (v/v) neutral buffered formalin (Sigma #HT501128-4L) for 24 h and then transferred to 70% ethanol.

### Histology

Processing, paraffin embedding, sectioning, and hematoxylin and eosin staining were performed by the Reference Histology Core Facility (Johns Hopkins University) according to standard protocols. Slides were scanned using Zeiss Axio Scan.Z1 slide scanner and ZEN Slidescan 2.3. Images were processed using QuPath (v.0.4.4).

### Cell Proliferation Assays

Cells were plated at 3,000 cells in 100 µL per well in 96-well plates. Medium was changed 24 h later according to indicated treatment. 20 µL of CellTiter 96 AQueous One Solution Cell Proliferation Assay (Promega #G3582) reagent was added to each well upon completion of experiment duration. Plates were incubated in 37°C, 5% CO_2_ for 1-4 h. Absorbance at 490 nm was read using FLUOstar Omega microplate reader (BMG LABTECH) to quantify cell viability.

### Drug Synergy Assays

Cells were seeded in 384-well plates at 250 cells in 50 µL of either 10% LPDS or 10% FBS media supplemented with 1 mM mevalonate (MilliporeSigma #M4667), 5 µg/mL cholesterol (MilliporeSigma #C3045) in ethanol, and 20 µM oleic acid-albumin (MilliporeSigma #O3008). After 24 h, the indicated compounds were applied using the semi-automated Tecan D300e Digital Dispenser, normalized to 0.5% DMSO. Drug compounds were tested across a logarithmically scaled curve: Fluvastatin and Simvastatin (range: 2.6x10^-6^ to 4.0x10^-2^ M), and TMDP (range: 2.6x10^-6^ to 1.0x10^-2^ M). Cells were allowed to grow for 5 days. Cell viability was measured using alamarBlue^TM^ Cell Viability Reagent (ThermoFisher #DAL1100) per the manufacturer’s protocol. Plates were incubated with the reagent for 4 h and fluorescence measured using the FLUOstar Omega microplate reader (BMG LABTECH) with the following parameters: excitation at 544 nm, emission at 590 nm, gain at 1000. Data were analyzed and plotted for synergy using the web-based tool SynergyFinder 2.0 (20). Curve fitting was achieved using a four-parameter logistic regression algorithm, and synergy calculated using the zero-interaction potency (ZIP) reference model (21).

### Apoptosis Assays

Cells were plated in 6-well plates at 1.5 x 10^5^ cells per 2 mL. Medium was changed 24-48 h later according to indicated treatment. Apoptosis was quantified using the Dead Cell Apoptosis Kit (Thermo #V35113) following the manufacturer’s protocol. Samples were filtered through 100 µm Nylon cell strainer (Falcon #352360) and stained for 15 min at room temperature. Apoptotic cell percentages were calculated by quantifying the number of Annexin V-FITC positive cells on the Attune NxT flow cytometer (ThermoFisher). Data were analyzed using FlowJo software (v.10.8.1).

### Real-time quantitative PCR

Total RNA was extracted using the Monarch Total RNA Miniprep Kit (New England Biolabs #T2010S). Reverse transcription was performed using iScript Reverse Transcription Supermix (BioRad #1708841) according to the manufacturer’s protocol. Gene expression was quantified using PowerUp SYBR Green Master Mix (ThermoFisher #A25741) on BioRad CFX Opus 96. *GAPDH* expression was used as an internal control to calculate relative gene expression. Primers are listed in **Table S1**.

### RNA-seq

Total RNA was extracted using the Monarch Total RNA Miniprep Kit (New England BioLabs #T2010S) and submitted to Azenta Life Sciences for library preparation and Illumina sequencing.

### Statistical analysis

Statistical analyses were performed using GraphPad Prism (v.10.2.0). Data are presented as described in corresponding figure legends.

## Results

### CRISPR screening identified terpenoid backbone biosynthesis as a PDAC tumor growth dependency

SREBPs are responsible for controlling a myriad of lipid metabolic genes and are required for PDAC tumor growth in vivo (17) **(Fig. 1A)**. Specifically, we previously demonstrated that *SCAP* is required for PDAC tumor growth and development. To probe the mechanism behind SREBP pathway essentiality in PDAC tumor cells, we constructed a focused CRISPR knockout library containing 10 sgRNAs per gene against 148 known SREBP target genes and 10 positive control genes, such as *KRAS*, *SREBP1*, *SREBP2*, and *SCAP*. The target gene list was derived from two previous studies identifying genes regulated by SREBPs in mouse liver by microarray and in human liver cells by ChIP-seq (22,23). Non-targeting sgRNAs were also included to establish a baseline for the screen. A complete list of genes and sgRNA sequences is provided in **Table S2.** Illumina sequencing confirmed that our plasmid library had a normal distribution of sgRNAs **(Fig. S1A)**. We then performed loss-of-function genetic screens using a patient-derived *KRAS^G12D^/TP53^L344P^* PDAC cell line (Pa03c) stably expressing Cas9 (24). To compare dependencies of 2D monolayer cell culture and tumors, we conducted in vitro and in vivo screens in parallel **(Fig. 1B)**.

**Figure 1.**
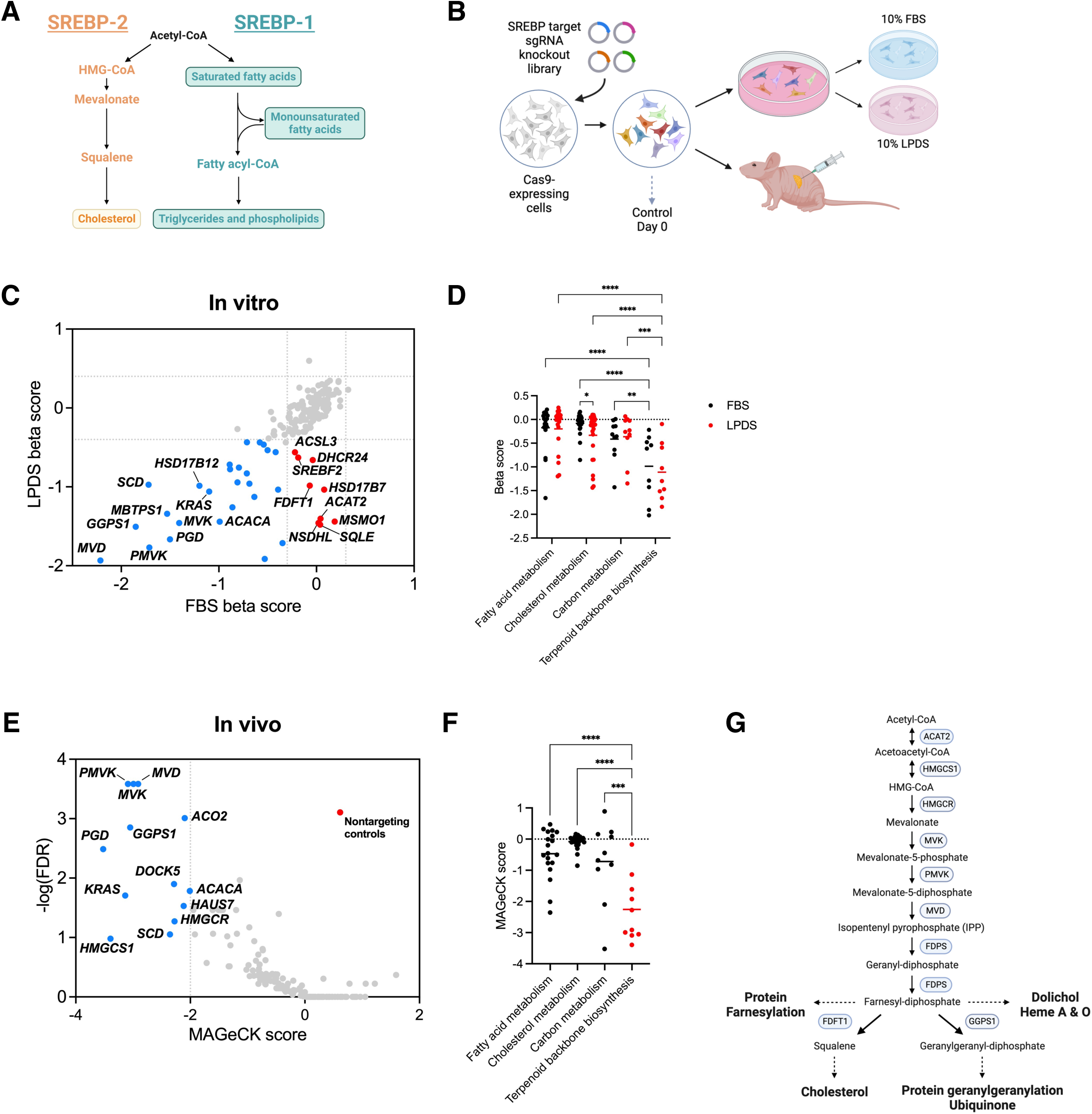
CRISPR screening identifies SREBP target gene growth dependencies in PDAC tumor cells. (A) SREBPs maintain lipid homeostasis by regulating lipogenic gene transcription. Shown are the two arms of the SREBP pathway. SREBP-1 primarily controls fatty acid synthesis and SREBP-2 primarily controls cholesterol synthesis. (B) Cas9-expressing Pa03c cells were infected with an SREBP target gene focused CRISPR knockout viral library. Following selection, cells were collected for a reference sample and then either cultured in 10% FBS or 10% LPDS or injected into the pancreas of nude mice. (C) In vitro CRISPR screening beta scores for FBS and LPDS conditions after 12 days. Blue dots show genes with beta scores < -0.4 in both FBS and LPDS conditions; red dots show genes with beta scores < -0.4 in LPDS condition only. A negative beta score indicates depletion, while a positive score indicates enrichment, relative to the day 0 control sample. (D) Beta scores from in vitro screens in FBS (black) and LPDS (red) for genes belonging to fatty acid metabolism (n=19), cholesterol metabolism (n=26), carbon metabolism (n=10), and terpenoid backbone biosynthesis (n=9) KEGG pathways. Data are shown as mean with each point representing one gene, Two-way ANOVA, *p <0.05, **p <0.01, ***p<0.001, ****p<0.0001. (E) In vivo CRISPR screen MAGeCK scores vs. -logFDR (n = 7 independent tumors). Blue dots show genes with MAGeCK scores < - 2; red dot shows negative control guides. (F) MAGeCK scores from in vivo screen for genes belonging to fatty acid metabolism, cholesterol metabolism, carbon metabolism, and terpenoid backbone biosynthesis KEGG pathways. Means are shown, One-way ANOVA, ***p<0.001, ****p<0.0001. (G) The mevalonate pathway synthesizes cholesterol and non-sterol isoprenoids such as dolichol, heme, farnesyl-diphosphate, geranylgeranyl diphosphate, and ubiquinone.

For the in vitro screen, library-infected cells were grown in medium containing either 10% fetal bovine serum (FBS) or 10% lipoprotein-deficient serum (LPDS) **(Fig. 1C)**. We detected >99% of sgRNAs in each sample, indicating sufficient library representation **(Fig. S1B)**. Using MAGeCK, we calculated beta scores to measure gene selection, where positive scores indicate positively selected genes, and negative scores indicate genes negatively selected. To determine which metabolic pathways were essential for growth, we classified genes based on their KEGG pathway annotation and compared the beta scores of fatty acid metabolism, cholesterol metabolism, carbohydrate metabolism, and terpenoid backbone biosynthesis genes between the FBS and LPDS conditions **(Fig. 1D)**. When comparing beta scores between FBS and LPDS within each group, only cholesterol metabolism showed differential essentiality in the LPDS condition. This is to be expected, as the major difference between the two conditions is the presence of lipoprotein, a primary source of cholesterol. Interestingly, there was little to no requirement for fatty acid synthesis as demonstrated by average beta scores near zero in both conditions **(Fig. 1D)**. Terpenoid backbone biosynthesis genes, such as *GGPS1, MVK, PMVK,* and *MVD,* in both FBS and LPDS, showed the lowest beta scores when compared to every other gene set.

For the in vivo screen, we injected library-infected cells into the pancreata of nude mice. Tumors (n=7) were collected after 4 weeks of growth and were treated as independent samples **(Fig. 1E)**. Due to the small size of the custom library (∼2500 sgRNAs), we were able to detect ∼90% of sgRNAs in each tumor despite the low engraftment efficiency of the orthotopic xenograft model **(Fig. S1C)**. As observed in the in vitro screens, terpenoid backbone biosynthesis pathway gene scores were significantly lower than the other gene sets **(Fig. 1F)**. Surprisingly, there was no statistical difference between any other two gene sets, suggesting a lesser requirement for cholesterol and fatty acid synthesis in PDAC tumor growth in vivo. As expected, we observed positive control *KRAS* as an essential gene in both screens. Notably, many mevalonate pathway genes showed similar or lower scores than *KRAS*. Comparing the two screening approaches, results from the in vivo screen more closely resembled the results from the FBS condition of the in vitro screen, as indicated by linear regression comparison of gene scores (FBS R^2^ = 0.57; LPDS R^2^ = 0.45) **(Fig. S1D)**. A full list of gene scores and KEGG pathway annotations are provided in **Table S3**.

Taken together, our CRISPR screens indicate that the terpenoid backbone biosynthesis genes are essential for PDAC tumor cell growth in vitro and in vivo. These genes, collectively referred to as the mevalonate pathway, support the synthesis of isoprenoids necessary for farnesyl, geranylgeranyl, heme, ubiquinone, cholesterol, and dolichol production and have been identified previously as a potential therapeutic target in cancer (25) **(Fig. 1G)**. Despite including genes required for cholesterol synthesis, we did not observe essentiality for these, indicating that the mevalonate pathway is providing an essential metabolite other than cholesterol. These data also suggest that tumor cells acquire cholesterol and fatty acids from means other than de novo biosynthesis. The mevalonate pathway was therefore prioritized for further study to understand the mechanism behind this requirement.

### Mevalonate pathway activity is essential for PDAC cell growth

To validate the results from our CRISPR screens and determine the therapeutic efficacy of targeting the mevalonate pathway in PDAC, we generated a dox-inducible CRISPRi *HMGCR* knockdown cell line **(Fig. 2A)**. *HMGCR* catalyzes the formation of mevalonate and is commonly targeted pharmacologically by statins. Using an orthotopic xenograft model in nude mice, we observed that *HMGCR* knockdown tumors were significantly reduced in size compared to the untreated group **(Fig. 2B)**. Out of 8 animals, 6 did not develop resectable pancreatic tumors in the dox-treated group, compared to only 1 in the untreated group. H&E tissue staining confirmed the absence of tumor lesions in the pancreata extracted from the mice without visible tumor growth **(Fig. 2C)**. Histologically, the pancreas appeared architecturally normal in the doxycycline chow-fed animals that did not develop tumors, while tumors in the untreated group showed characteristic neoplastic cell morphology that was consistent with the parent cell line. We also engineered a nontargeting control cell line to control for the effect of doxycycline on tumor growth **(Fig. S2A)**. There was no significant difference in tumor size or histologic analysis between the normal chow and doxycycline chow groups **(Fig. S2B-C).**

**Figure 2.**
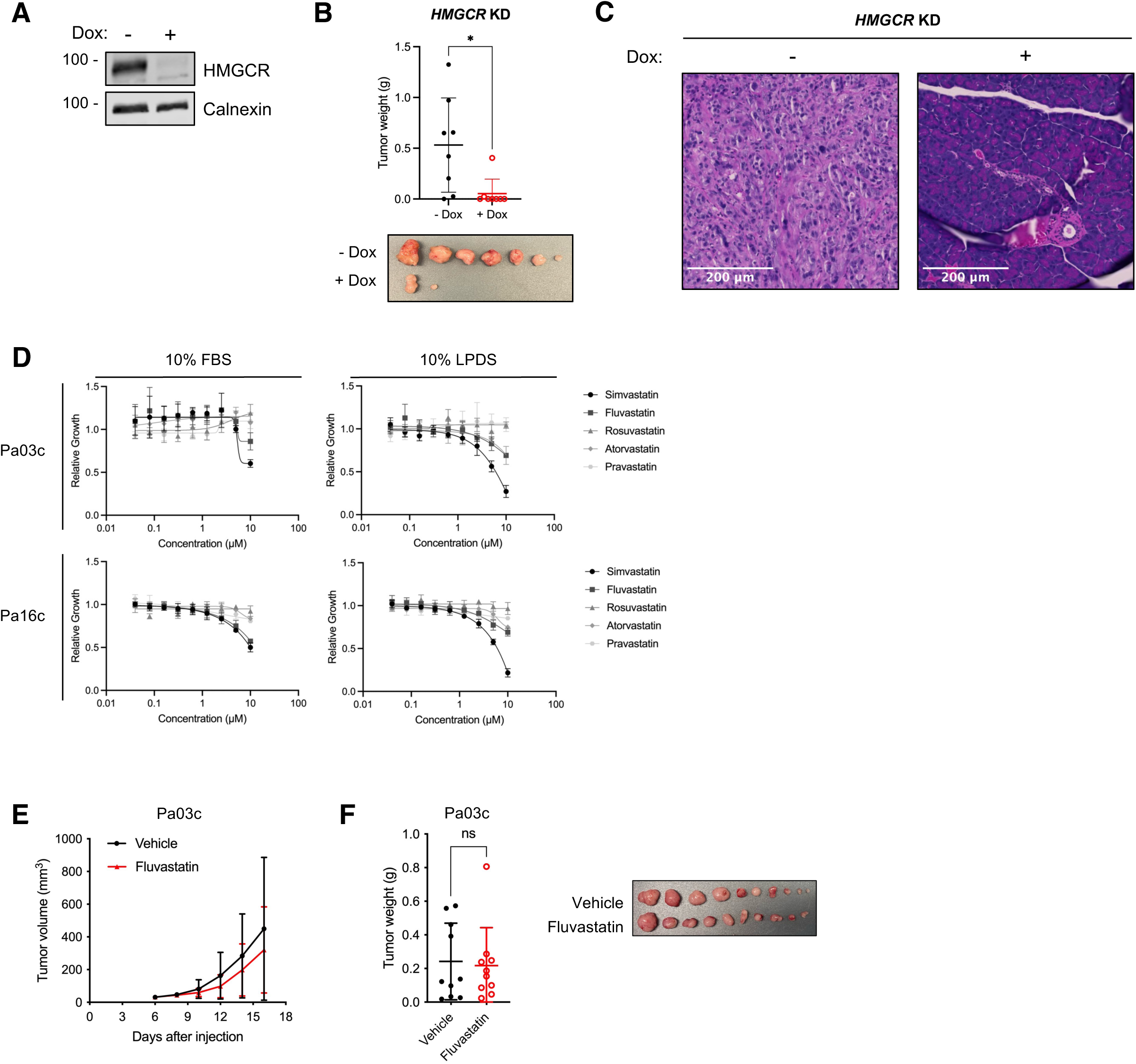
Mevalonate pathway activity is essential for PDAC tumor growth. (A) Western blots of Pa03c *HMGCR* knockdown monoclonal cell line treated without or with 1 µg/ml doxycycline as indicated for 72 h. (B) Orthotopic xenograft tumor weights of dox-inducible *HMGCR* knockdown cells after 1 week of growth followed by 3 weeks of normal chow or doxycycline treatment (n = 8 mice per group, mean ± SD, unpaired t-test, *p<0.05). Individual tumors are shown. (C) H&E images of tumors within pancreata in (B) from normal chow group (left) or from doxycycline chow group (right). (D) Growth curves of Pa03c and Pa16c cells treated with 10% FBS or 10% LPDS and varying concentrations of statins for 72 h (n = 3 biological replicates, mean ± SD). (E) Pa03c subcutaneous xenograft tumor volumes from mice treated orally with vehicle or 50 mg/kg Fluvastatin daily (n = 10 mice per group, mean ± SD). (F) Tumor weights on day 16 from vehicle- or Fluvastatin-treated mice shown in (E) (n = 10 mice per group, mean ± SD, unpaired t-test, ns=not significant). Individual tumors are shown.

Since statins are commonly prescribed FDA-approved drugs that inhibit HMGCR, we next measured cell growth of multiple patient derived PDAC cell lines across a panel of statins commonly used in clinical settings. Notably, we observed varying responses, with more effective growth inhibition in 10% LPDS, perhaps due to limited supply of extracellular cholesterol **(Fig. 2D, Fig. S2D)**. To test statin efficacy on PDAC tumor growth, we used a subcutaneous xenograft model in nude mice. Pa03c tumor size was measured every other day to assess growth over time **(Fig. 2E)**. We observed no significant difference in tumor weight between groups, suggesting that statins alone do not affect tumor growth in vivo **(Fig. 2F)**. Similar results were also observed in Pa16c cells **(Fig. S2E-F)**. Given the tumor suppressive in vivo results with CRISPRi knockdown of *HMGCR,* the in vivo statin results were discordant.

### Statins synergize with SREBP inhibitors to induce apoptosis

Despite the large body of literature suggesting that statins may be beneficial for cancer therapy, statins have yet to be approved as a cancer treatment due to wide variations in response and dose-limiting toxicity (26). Moreover, association studies have concluded that statin use is not associated with reduced risk of pancreatic cancer (27). As one possible explanation, we hypothesized that statin treatment limits cholesterol synthesis, which triggers SREBP activation and results in further upregulation of *HMGCR* (and other mevalonate pathway genes). Therefore, we tested if cell death could be amplified by treatment with an SREBP inhibitor to prevent this homeostatic response **(Fig. 3A)**. Consistent with this hypothesis, previous studies in prostate cancer and acute myeloid leukemia have shown that SREBP inhibitor and statin combination treatments are more effective than monotherapy (28,29). These studies utilized dipyridamole as an SREBP inhibitor, but we instead opted to test its more specific derivative, N2,N2,N6,N6-tetrakis[2-methoxyethyl]-4,8-di[piperidin-1-yl]pyrimido[5,4-d]pyrimidine-2,6-diamine (TMDP) (30). Upon treating cells with TMDP, we observed lipid-dependent growth inhibition, as expected, due to SREBP activity being essential under low lipid conditions **(Fig. 3B, Fig. S3A)**. To test our hypothesis that the SREBP gene response is activated upon statin treatment, we assessed *HMGCR*, *HMGCS1*, and *INSIG1* target gene expression by qPCR **(Fig. 3C)**. As predicted, Fluvastatin treatment upregulated these SREBP target genes, and TMDP inhibited baseline gene expression in 10% LPDS. Importantly, TMDP inhibited statin-dependent upregulation of SREBP target genes to levels approaching that with TMDP alone.

**Figure 3.**
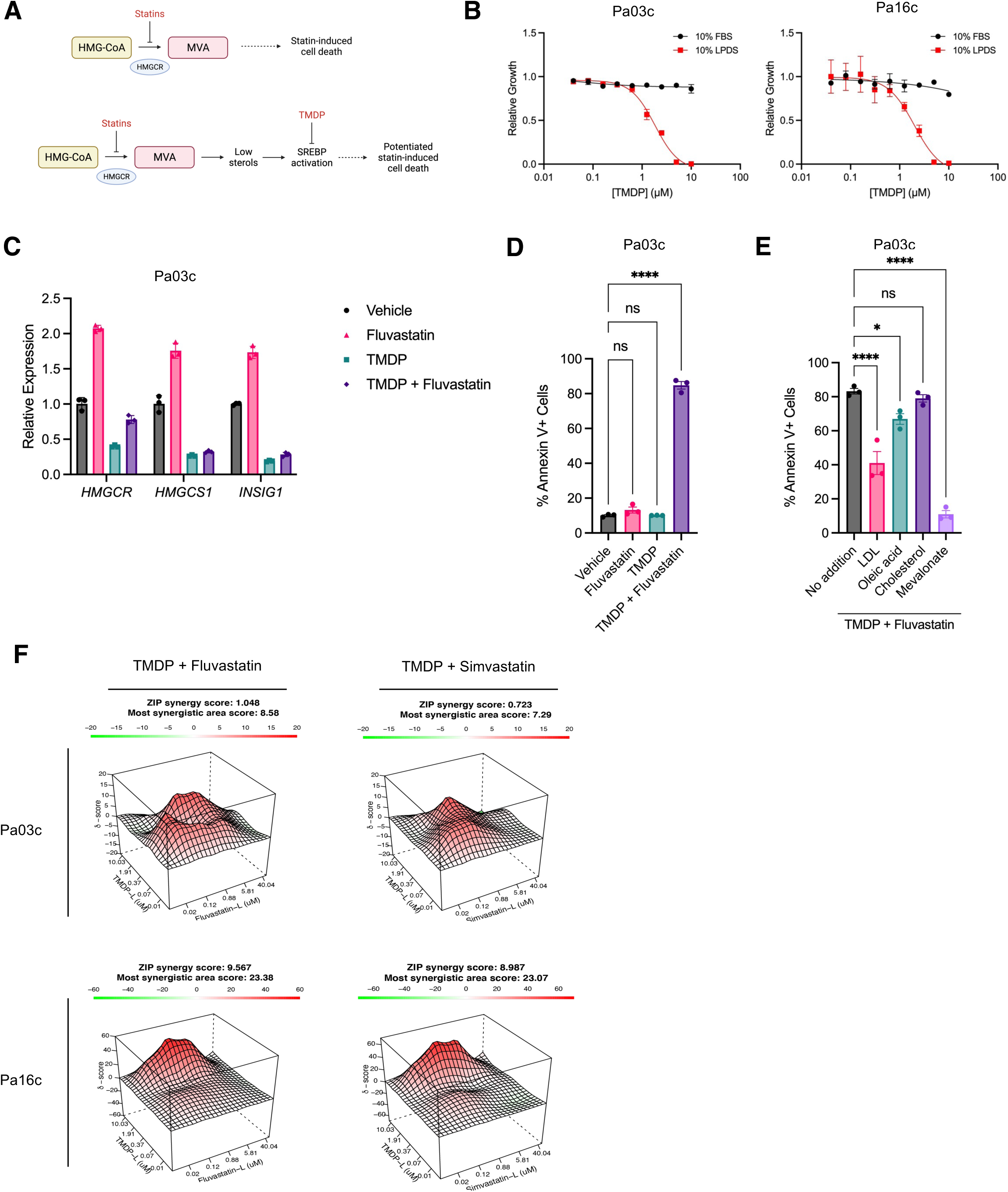
Statins induce apoptosis synergistically with SREBP inhibitors in vitro. (A) Schematic outlining rationale for combination drug treatment. (B) Growth curves of Pa03c and Pa16c cells cultured in 10% FBS or 10% LPDS and varying concentrations of TMDP for 72 h (n = 3 replicates, mean ± SD). (C) Quantitative PCR of *HMGCR, HMGCS1,* and *INSIG1* in Pa03c cells cultured in 2.5 µM TMDP and/or 5 µM Fluvastatin in 10% LPDS for 24 h (n = 3 technical replicates, mean ± SD). (D) Annexin V flow cytometry quantification of Pa03c cells cultured in 2.5 µM TMDP and/or 5 µM Fluvastatin in 10% LPDS for 48 h (n = 3 biological replicates, mean ± SD, One-way ANOVA, ****p<0.0001). (E) Pa03c cells treated with combination of TMDP and Fluvastatin as in (D) with the addition of 10 µg/mL LDL, 20 µM oleic acid, 5 µg/mL cholesterol, or 1 mM mevalonate (n = 3 biological replicates, mean ± SD, One-way ANOVA, *p<0.05, ****p<0.0001). (F) ZIP synergy score plots for TMDP + Fluvastatin or TMDP + Simvastatin in Pa03c and Pa16c cells (n = 2 biological replicates for each dose). Each map shows the synergistic (red) and antagonistic (green) dose regions. The most synergistic area score is the average score of the highest 3x3 dose-window average across the entire matrix. A positive score indicates synergy while a negative score indicates antagonism.

To examine the effect of these treatments on cell viability, we assayed Annexin V staining by flow cytometry. While there was no significant change in apoptosis when the cells were treated with Fluvastatin or TMDP alone, we observed a stark increase in apoptosis when PDAC cells were treated with the drug combination **(Fig. 3D)**. These results were also observed with a separate SREBP pathway inhibitor, PF-429242, to confirm that this effect was not specific to TMDP **(Fig. S3B)**. SREBPs control cellular lipid supply, so we next examined which lipid becomes limiting during treatment with Fluvastatin and TMDP. Given that SREBPs supply fatty acids and cholesterol to cells **(Fig. 1A)**, we tested low-density lipoprotein (LDL), oleic acid, cholesterol, and mevalonate. We observed that LDL and oleic acid were able to rescue cell death to some extent, but only mevalonate fully rescued, returning apoptosis to basal levels **(Fig. 3E)**. These data indicate that mevalonate provides an essential metabolite for cell survival separate from cholesterol and that fatty acids can partially rescue this cell death. Importantly, we established that restricting mevalonate synthesis through simultaneous SREBP and HMGCR inhibition results in synergistic cell death. These results corroborate our in vivo genetic knockout screen that illustrates the essentiality of mevalonate pathway activity for tumor growth, independent from cholesterol synthesis.

To determine the broader synergy profile of TMDP and statins, we assessed cell growth across a range of drug concentrations for cells treated with combinations of TMDP and either Fluvastatin or Simvastatin **(Fig. 3F, S3C)**. We calculated zero interaction potency (ZIP) synergy scores that indicate the average of all scores across the entire landscape (21). There were several drug combinations that scored as synergistic in each landscape, indicating that these drugs act synergistically across multiple cell lines to impair PDAC cell growth.

### Bulk RNA-seq reveals broad cellular stress in response to combination drug treatment

Treating PDAC cells with TMDP and Fluvastatin induces synergistic apoptotic cell death. To better understand the mechanism underlying induction of apoptosis, we conducted bulk RNA-seq using cells treated with each drug individually or in combination. We performed gene set enrichment analysis (GSEA) using the Hallmark gene set to analyze each condition and statistically significantly enriched pathways (FDR < 0.1) were plotted **(Fig. 4A)**. In agreement with our hypothesis that SREBP activation occurs upon Fluvastatin treatment, we observed that while fatty acid metabolism and cholesterol homeostasis gene sets were enriched in the Fluvastatin condition, these gene sets were depleted in the TMDP treatment condition **(Fig. 4B-C)**. Moreover, in the combination treatment, these gene sets were no longer enriched significantly, indicating that TMDP was able to blunt the lipid homeostatic response induced by Fluvastatin, as illustrated in the NES plots **(Fig. 4B)**. Additionally, we observed that cell cycle genes were depleted in the drug combination treatment, including G2M checkpoint genes and E2F targets. We conducted a parallel experiment using the SREBP pathway inhibitor PF-429242 and likewise observed that fatty acid metabolism and cholesterol homeostasis were downregulated in the combination treatment condition compared to Fluvastatin alone **(Fig. S4A-B)**.

**Figure 4.**
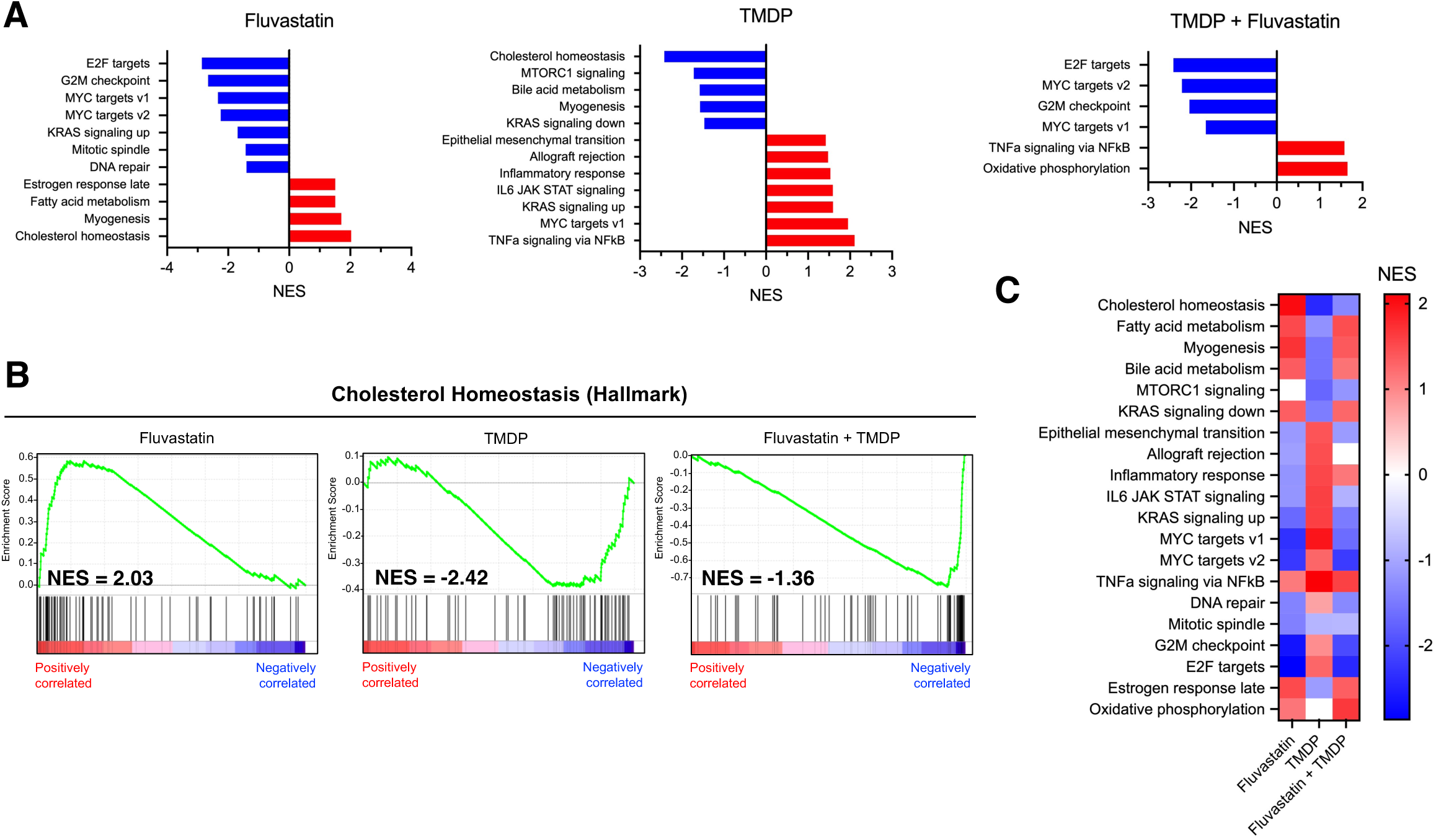
Bulk RNA-seq reveals broad cellular stress in response to combination drug treatment. (A) NES scores for Hallmark gene sets with FDR < 0.1 of RNA-seq samples from cells cultured in 5 µM Fluvastatin, 2.5 µM TMDP, or the drug combination in 10% LPDS for 16 h. (B) NES plots for Hallmark cholesterol homeostasis gene set for Fluvastatin (left), TMDP (middle), and Fluvastatin + TMDP (right) samples. (C) Heat map showing NES scores for all Hallmark gene sets identified in treated cells from (A).

When we plotted the NES scores of all these gene sets across the 3 conditions, we observed that the combination treatment mimics the signature of Fluvastatin most closely, apart from cholesterol homeostasis, MTORC1 signaling, and inflammatory response **(Fig. 4C)**. We also observed that these expression changes were similar when we added mevalonate to the medium, but without the inflammatory response or other stress response pathways such as TNF-α signaling, suggesting that a lack of mevalonate induces these stress responses **(Fig. S4C)**. Together, these data confirm that our combination drug treatment can dampen lipid homeostatic responses and induce stress responses within PDAC cells.

### Mevalonate pathway inhibition results in protein prenylation defects due to GGPP depletion

We next were interested in understanding which mevalonate pathway product was essential for PDAC cell survival. Farnesyl-diphosphate (FPP) and geranylgeranyl diphosphate (GGPP) are two isoprenoid mevalonate pathway products that act as post-translational modifications of proteins and support dolichol, ubiquinone, and heme synthesis **(Fig. 1G)**. Importantly, many small GTPases such as RAS are isoprenylated and require such modifications to function and localize appropriately (31). We tested a geranylgeranyl transferase inhibitor (GGTI-298) and a farnesyl transferase inhibitor (FTI-277) to determine if inhibiting protein isoprenylation had effects on cell growth in vitro **(Fig. 5A)**. GGTI-298 showed growth suppression at higher doses while FTI-277 had little effect on growth in both media conditions up to 10 µM. To determine if protein prenylation can be restricted to result in synergistic apoptosis, we tested synergy between TMDP and GGTI-298 or FTI-277. We also tested synergy with a squalene epoxidase inhibitor (YM-53601) to determine if cholesterol synthesis inhibition could also induce synergistic cell death. We observed that GGTI-298 and FTI-277 were able to synergize with TMDP to induce apoptosis but to a lesser extent than Fluvastatin **(Fig. 5B)**. YM-53601 did not synergize with TMDP, further supporting the conclusion that cholesterol is not the limiting metabolite in these cells. Notably, cell death was completely rescued by the addition of GGPP upon Fluvastatin-TMDP combination drug treatment, suggesting that cell death is a result of GGPP starvation **(Fig. 5C)**.

**Figure 5.**
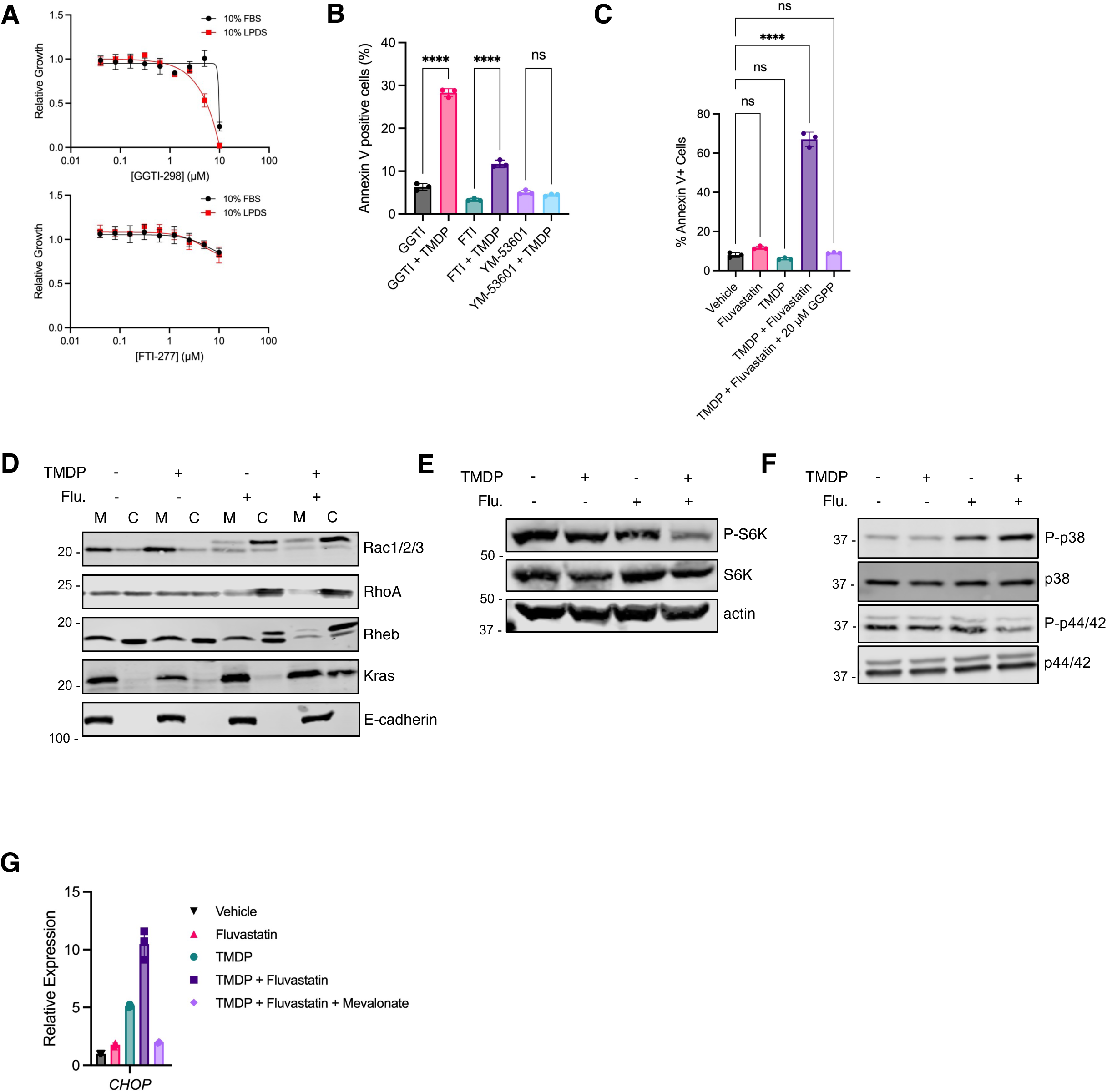
Mevalonate pathway inhibition results in protein prenylation defects due to GGPP depletion. (A) Growth curves of Pa03c cells cultured in 10% FBS or 10% LPDS and indicated concentrations of GGTI-298 or FTI-277 for 72 h (n = 3 replicates, mean ± SD). (B) Annexin V staining of Pa03c cells cultured in 2.5 µM TMDP and/or 5 µM GGTI-298, 5 µM FTI-277, 5 µM YM-53601 in 10% LPDS for 48 h (n = 3 biological replicates, mean ± SD, One-way ANOVA, ****p<0.0001, ns=not significant). (C) Annexin V staining of Pa03c cells cultured in 10% LPDS with 5 µM Fluvastatin, 2.5 µM TMDP, or the drug combination without or with 20 µM GGPP for 48 h (n = 3 biological replicates, mean ± SD, One-way ANOVA, ****p<0.0001, ns=not significant). (D) Western blots of membrane (M) and cytosolic (C) protein fractions from Pa03c cells cultured in 2.5 µM TMDP, 5 µM Fluvastatin, or the drug combination in 10% LPDS for 24 h. (E) Western blots assaying total S6 Kinase and phospho-S6 Kinase levels in Pa03c cells cultured in 2.5 µM TMDP, 5 µM Fluvastatin, or the drug combination in 10% LPDS for 24 h. (F) Western blots assaying MAPK proteins p38 and p44/42 total and phosphorylated levels in Pa03c cells cultured in 2.5 µM TMDP, 5 µM Fluvastatin, or the drug combination in 10% LPDS for 24 h. (G) Expression levels of ER stress marker *CHOP* in Pa03c cells cultured in 10% LPDS with 5 µM Fluvastatin, 2.5 µM TMDP, or the drug combination without or with 1 mM mevalonate for 30 h (n = 3 technical replicates representative of two independent experiments, mean ± SD).

GGPP acts primarily as a post-translational modification, suggesting that defects in protein prenylation may contribute to the mechanism behind the observed cell death. To test this, we evaluated the localization and prenylation status of several small G proteins by cell fractionation. Treatment with TMDP alone had no effect either prenylation or protein localization **(Fig. 5D)**. For multiple small G proteins, we observed an increase in unprenylated proteins (upper bands) when cells were treated with Fluvastatin as well as a shift into the cytosolic protein fraction. Consistent with the role of SREBP in the adaptive response to Fluvastatin treatment, these effects were exacerbated when cells were treated with the combination of inhibitors. Strikingly, KRAS localization was unchanged when treated with Fluvastatin or TMDP alone, but there was a notable shift into the cytosolic fraction in the combination treatment condition. This was also the most dramatic change observed across small G proteins tested.

Considering the important role that small G proteins play in signaling networks, we chose to test the effect of the combination drug treatment on MTORC1 and MAPK signaling pathway activity, since these are activated downstream of both *RHEB* and *KRAS*, respectively. We observed a reduction in S6 kinase phosphorylation as a marker of MTORC1 activation in the combination treatment but not in either single drug treatment **(Fig. 5E)**. Additionally, we observed an increase in phosphorylated p38, signaling an activation in the stress response, and a modest decrease in phosphorylated p44/42, representing reduced proliferative signaling **(Fig. 5F)**. We also observed upregulation of *CHOP* expression, indicating that the ER stress response is activated upon combination drug treatment, but was reduced with mevalonate addition **(Fig. 5G)**. Together, these data illustrate that mevalonate pathway inhibition induces pleotropic effects on protein localization and cellular signaling. Importantly, these results are consistent with those observed in bulk RNA-seq and reveal the functional consequences of limiting isoprenoid synthesis.

### GGPS1 is essential for PDAC tumor growth

Geranylgeranyl diphosphate synthase (GGPS1) has been recognized previously as a potential anticancer target, illustrating that inhibiting GGPP synthesis can perturb cancer cell growth (32). Our results align with this idea and indicate that PDAC tumors require mevalonate pathway activity to support GGPP synthesis. Therefore, we chose to investigate *GGPS1* directly. Examining patient pancreatic cancer samples from matched normal and tumor samples showed that *GGPS1* is significantly upregulated in tumors compared to normal tissue **(Fig. 6A)** (33). We then generated *GGPS1* dox-inducible knockdown cell lines to test the requirement for tumor growth in vivo **(Fig. 6B)**. After 3 weeks of doxycycline chow treatment, we observed that tumors were significantly smaller as reflected by tumor weight after resection **(Fig. 6C)**. These in vivo results further validate our CRISPR screening results that show a requirement for the mevalonate pathway and our conclusion that GGPP is essential for PDAC tumor growth.

**Figure 6.**
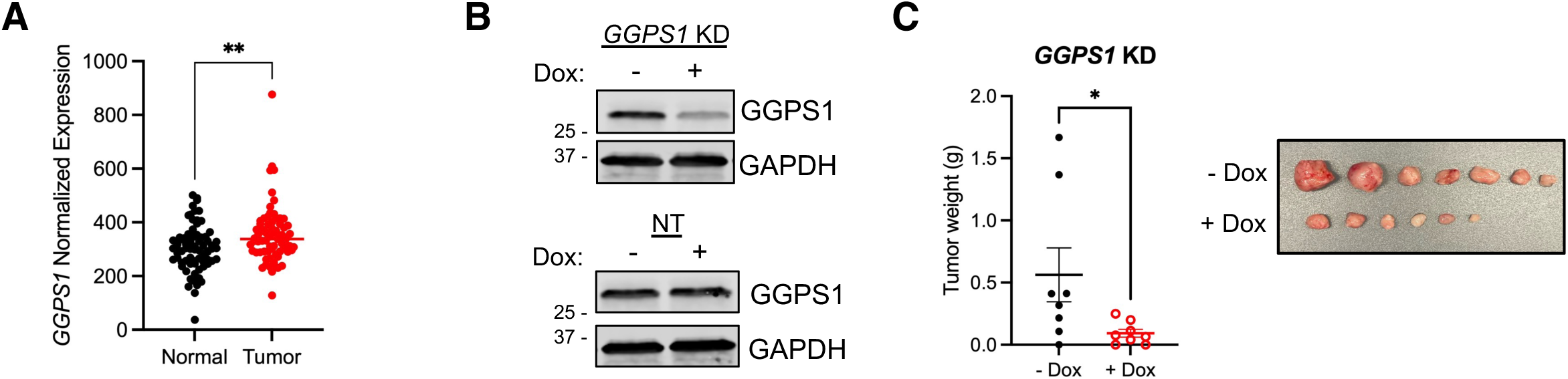
GGPS1 is essential for PDAC tumor growth. (A) TN-plot gene chip data showing *GGPS1* expression in normal pancreas tissues and tumor tissues (unpaired t-test, **p<0.01). (B) Western blot of GGPS1 and GAPDH for dox-inducible non-targeting (NT) and *GGPS1* knockdown (KD) cell lines treated without or with 1 µg/ml doxycycline as indicated for 72 h. (C) Orthotopic xenograft tumor weights of dox-inducible *GGPS1* knockdown cells after 1 week of growth followed by 3 weeks of normal chow or doxycycline treatment (n = 8 mice per group, mean ± SD, unpaired t-test, *p<0.05). Individual tumors are shown.

## Discussion

SREBPs are central in maintaining lipid homeostasis, and it has been well established that lipid metabolism is a critical facet of metabolic reprogramming in cancer (2,5). Our recent work illustrated that *SCAP* is required for tumor growth in orthotopic and subcutaneous xenografts (17). In this study, we dissected the requirement of SREBPs in PDAC tumor growth using genetic knockout screens. We concluded that the mevalonate pathway is essential for tumor establishment in an orthotopic xenograft mouse model, and that this requirement is based on the need for GGPP. Taken together, these results further support the pursuit of SREBPs as a target and establish a mechanism underlying the overarching essentiality of SREBPs in PDAC.

Our results extend previous knowledge demonstrating the essentiality of the mevalonate pathway in various cancer types, which has been reviewed elsewhere (25,34). Adding to this body of literature, we contribute a mechanistic study on the specific isoprenoid requirement of the mevalonate pathway for tumor growth in PDAC that ties together lipid metabolism and cellular oncogenic signaling. Specifically, we show that there are functional downstream consequences on signaling due to defects in protein prenylation. Moreover, we demonstrate that synergistic apoptosis can be achieved when PDAC cells are treated simultaneously with a statin and SREBP inhibitor, and that this effect is GGPP-dependent. Excitingly, we also observed a stark synergistic response in KRAS localization in addition to other small G proteins tested. We propose that this combination drug treatment works more effectively than statins alone due to the SREBP-dependent homeostatic response induced upon statin treatment. This is also a potential explanation why genetic knockdown of *HMGCR* was effective in suppressing tumor growth, but statins alone were not. Despite the lack of a specific bioavailable SREBP inhibitor to test in vivo, we present the mechanism behind a combination drug treatment that may serve as a solution to dose-limiting toxicity concerns of existing mevalonate pathway inhibitors like statins (26).

Altogether, we narrow down the requirement of the mevalonate pathway to GGPP. We showed that limiting GGPP synthesis through genetic knockdown of *GGPS1* restricted tumor growth in an orthotopic xenograft model. These data parallel the work done in previous studies exploring GGPS1 inhibitors as potential chemotherapeutic agents (32). It is important to note that aside from acting as a post-translational modification, GGPP also is required for ubiquinone synthesis, which was not explored in this study. It is possible that mitochondria are affected by a lack of ubiquinone production induced upon the combination drug treatment, which can result in oxidative stress and electron transport chain defects, as shown previously (35). Notably, we were not able to achieve the same level of synergy with the prenyl transferase inhibitors as with statins, indicating that a lack of ubiquinone could contribute to the observed phenotype. Supporting this idea, we observed that genes related to oxidative phosphorylation were upregulated in the combination drug treatment condition in our RNA-seq experiments **(Fig. 4A)**. Together, these data suggest that this synergistic effect may not be solely due to a lack of protein prenylation. A limitation of the present study is that our CRISPR knockout screen and other mouse studies were conducted in an immunodeficient model. This is an important consideration for future studies in determining the role of the mevalonate pathway in tumor development, owing to the fact that immune cell populations play a major role in tumor biology (36).

Overall, we define the mechanism behind the requirement of SREBPs in PDAC. Our results show that GGPP is an essential metabolite for tumor growth and that restricting GGPP synthesis genetically was sufficient to restrict tumor size in vivo. However, we currently lack a complete analysis of the effects of SREBP inhibition on established tumors and how these changes affect lipid species within tumors. Importantly, the present study reinforces the idea that SREBPs are an attractive target in PDAC and prompt further studies to identify a bioavailable pathway-specific inhibitor to be tested in vivo. Development of such a drug will better evaluate the therapeutic potential of targeting the SREBPs in cancer.

## Acknowledgements

We thank Joshua Laffin (Department of Cell Biology, Johns Hopkins School of Medicine) for generating and providing the Cas9-expressing cell line used for the genetic knockout screens. We also thank Timothy Osborne (Johns Hopkins All Children’s Hospital) for generously providing TMDP.

## Author Contributions

**C.S. Kubota:** Conceptualization, formal analysis, investigation, methodology, project administration, validation, visualization, writing—original draft, writing—review and editing; **S.L. Myers:** Conceptualization, formal analysis, funding acquisition, investigation, methodology, project administration, visualization, writing—review and editing; **T.T. Seppälä:** Methodology, resources, writing—review and editing; **R.A. Burkhart:** Project administration, resources, supervision, writing—review and editing; **P.J. Espenshade:** Conceptualization, funding acquisition, project administration, supervision, writing—review and editing.

## Data Availability

CRISPR knockout screening datasets generated during this study are available from the corresponding author on reasonable request. RNA-sequencing datasets generated during this study are available in the NCBI Gene Expression Omnibus (GEO) under accession number GSE247762.

## Tables

**Table S1.** Primer sequences.

**Table S2.** CRISPR knockout library sgRNA sequences

**Table S3.** CRISPR screen gene scores.

