## Supplementary Information for "In vivo CRISPR screening identifies geranylgeranyl diphosphate as a pancreatic cancer tumor growth dependency"

**Competing Interests:** T.T.S. reports a consultation fee from Amgen Finland, being a co-owner and CEO of Healthfund Finland Ltd, and member of the Clinical Advisory Board and minor stakeholder of LS Cancer Diag Ltd.

### Supplementary Figures

**A**

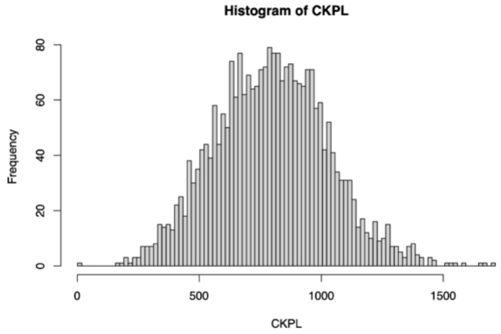


**B**

| | Label | Reads | Mapped | Percentage |
|---|---|---|---|---|
| 1 | 0A | 15871382 | 1350971 | 0.09 |
| 2 | 0B | 13584174 | 7086710 | 0.52 |
| 3 | 0C | 13433114 | 6846445 | 0.51 |
| 4 | 6LA | 16148030 | 8200052 | 0.51 |
| 5 | 6LB | 10259363 | 5252200 | 0.51 |
| 6 | 6LC | 11147020 | 6277514 | 0.56 |
| 7 | 9FA | 9627498 | 4921844 | 0.51 |
| 8 | 9FB | 12425058 | 7024685 | 0.57 |
| 9 | 9FC | 10626140 | 6032997 | 0.57 |
| 10 | 9LA | 15046547 | 7641381 | 0.51 |
| 11 | 9LB | 12862217 | 7263943 | 0.56 |
| 12 | 9LC | 13228312 | 7506294 | 0.57 |
| 13 | 12FA | 11559320 | 6446162 | 0.56 |
| 14 | 12FB | 14548210 | 7449591 | 0.51 |
| 15 | 12FC | 11543395 | 6315749 | 0.55 |
| 16 | 12LA | 17824217 | 10033978 | 0.56 |
| 17 | 12LB | 14857352 | 8393325 | 0.56 |
| 18 | 12LC | 16174980 | 9013943 | 0.56 |
| 19 | 3FA | 4507807 | 2448703 | 0.54 |
| 20 | 3FB | 3277515 | 1687058 | 0.51 |
| 21 | 3FC | 4243943 | 2433635 | 0.57 |
| 22 | 3LA | 2519009 | 1442524 | 0.57 |
| 23 | 3LB | 3185020 | 1721405 | 0.54 |
| 24 | 3LC | 4336286 | 2451299 | 0.57 |
| 25 | 6FA | 3901896 | 2210625 | 0.57 |
| 26 | 6FB | 4400247 | 2500150 | 0.57 |
| 27 | 6FC | 3299417 | 1863959 | 0.56 |

Table 1: Summary of comparisons

| | Label | TotalsgRNA | ZeroCounts | GiniIndex |
|---|---|---|---|---|
| 1 | 0A | 2560 | 1 | 0.05 |
| 2 | 0B | 2560 | 1 | 0.03 |
| 3 | 0C | 2560 | 1 | 0.03 |
| 4 | 6LA | 2560 | 1 | 0.06 |
| 5 | 6LB | 2560 | 3 | 0.06 |
| 6 | 6LC | 2560 | 1 | 0.06 |
| 7 | 9FA | 2560 | 4 | 0.07 |
| 8 | 9FB | 2560 | 6 | 0.06 |
| 9 | 9FC | 2560 | 5 | 0.06 |
| 10 | 9LA | 2560 | 7 | 0.07 |
| 11 | 9LB | 2560 | 2 | 0.06 |
| 12 | 9LC | 2560 | 6 | 0.07 |
| 13 | 12FA | 2560 | 5 | 0.07 |
| 14 | 12FB | 2560 | 12 | 0.07 |
| 15 | 12FC | 2560 | 11 | 0.07 |
| 16 | 12LA | 2560 | 9 | 0.08 |
| 17 | 12LB | 2560 | 10 | 0.08 |
| 18 | 12LC | 2560 | 8 | 0.07 |
| 19 | 3FA | 2560 | 2 | 0.05 |
| 20 | 3FB | 2560 | 2 | 0.05 |
| 21 | 3FC | 2560 | 1 | 0.05 |
| 22 | 3LA | 2560 | 2 | 0.05 |
| 23 | 3LB | 2560 | 2 | 0.05 |
| 24 | 3LC | 2560 | 2 | 0.05 |
| 25 | 6FA | 2560 | 3 | 0.06 |
| 26 | 6FB | 2560 | 7 | 0.07 |
| 27 | 6FC | 2560 | 17 | 0.07 |

Table 2: Summary of comparisons

**C**

| | Label | Reads | Mapped | Percentage |
|---|---|---|---|---|
| 1 | 0A | 1203230 | 654150 | 0.54 |
| 2 | 0B | 1443575 | 778671 | 0.54 |
| 3 | 0C | 1242880 | 237352 | 0.19 |
| 4 | TA | 1701992 | 944908 | 0.56 |
| 5 | TB | 1840926 | 1078380 | 0.59 |
| 6 | TC | 1831420 | 1100686 | 0.60 |
| 7 | TD | 1832850 | 781419 | 0.43 |
| 8 | TE | 2137870 | 1138949 | 0.53 |
| 9 | TF | 2050034 | 1148588 | 0.56 |
| 10 | TG | 1455060 | 724332 | 0.50 |

Table 1: Summary of comparisons

| | Label | TotalsgRNA | ZeroCounts | GiniIndex |
|---|---|---|---|---|
| 1 | 0A | 2560 | 2 | 0.05 |
| 2 | 0B | 2560 | 1 | 0.05 |
| 3 | 0C | 2560 | 1 | 0.06 |
| 4 | TA | 2560 | 50 | 0.27 |
| 5 | TB | 2560 | 331 | 0.40 |
| 6 | TC | 2560 | 378 | 0.40 |
| 7 | TD | 2560 | 235 | 0.33 |
| 8 | TE | 2560 | 150 | 0.31 |
| 9 | TF | 2560 | 246 | 0.37 |
| 10 | TG | 2560 | 236 | 0.37 |

Table 2: Summary of comparisons

**D**

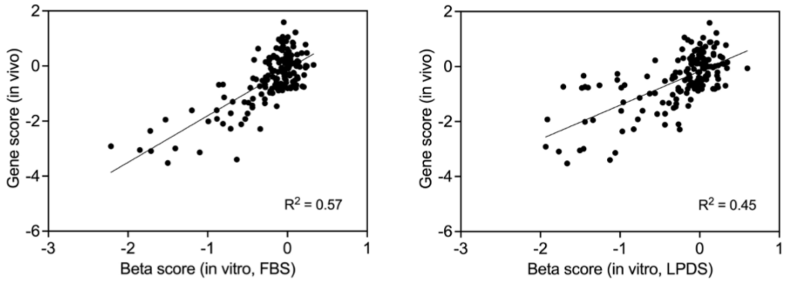


**Figure S1.** Custom CRISPR library screening. (A) Histogram of sgRNA distribution in custom CRISPR knockout plasmid library. (B) Tables showing number of reads, mapped reads, and zero counts from each condition of in vitro screen. Numbers in label indicate day; F=FBS, L=LPDS; A/B/C indicate biological replicates. (C) Tables showing number of reads, mapped reads, and zero counts from each mouse for in vivo screen. T indicates tumor; A/B/C/D/E/F/G indicates individual mouse sample. (D) In vivo CRISPR gene scores plotted against in vitro beta scores for (left) FBS, or (right) LPDS conditions. Linear regression $R^2$ value is shown.

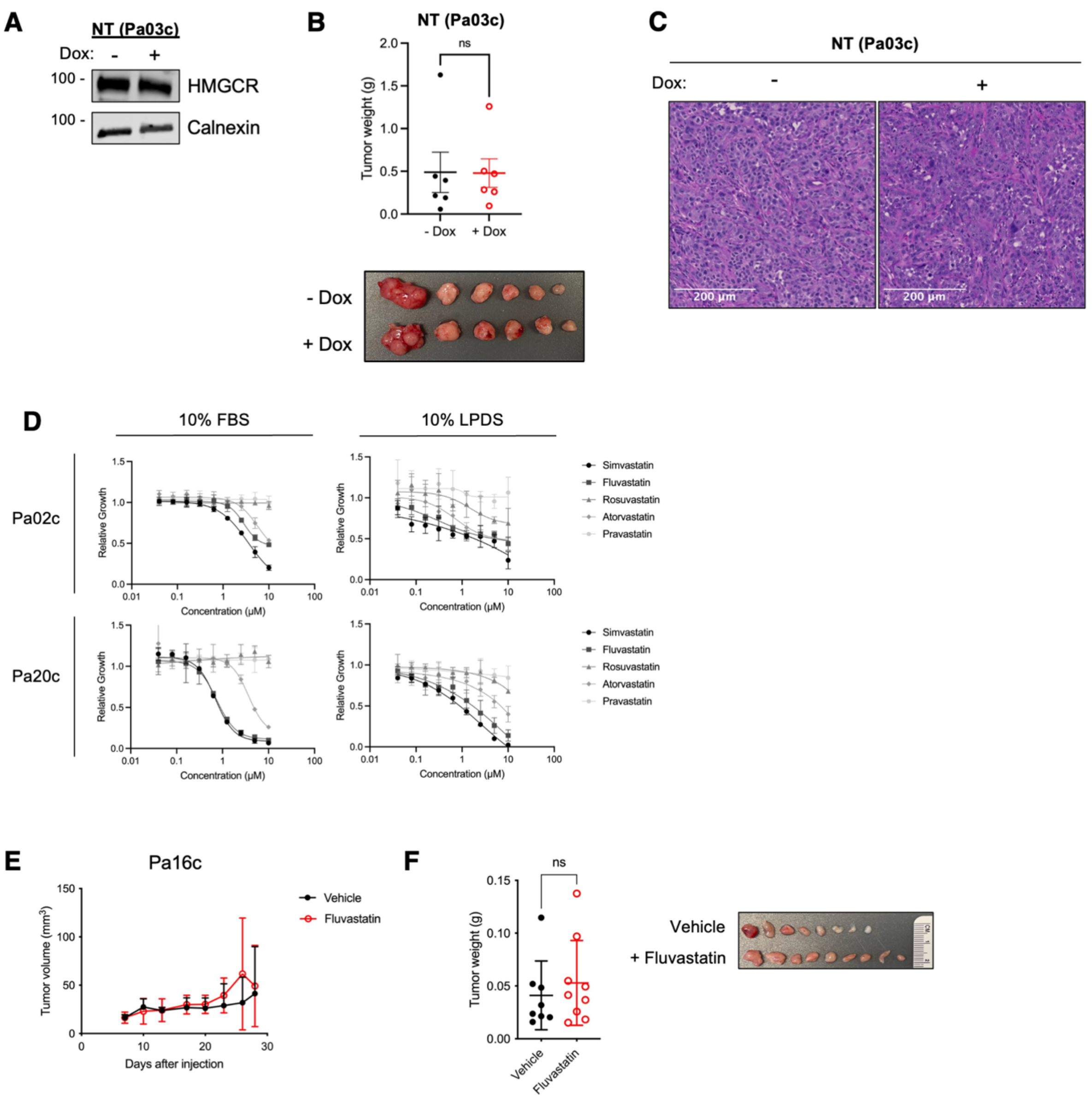


**Figure S2.** Mevalonate pathway activity is essential for PDAC tumor growth. (A) Western blots of dox-inducible non-targeting cell line treated without or with 1 μg/ml doxycycline as indicated for 72 h. (B) Orthotopic xenograft tumor weights of dox-inducible non-targeting knockdown Pa03c cells after 1 week of growth on normal chow followed by 3 weeks of either normal chow (-Dox) or doxycycline treatment (+Dox) (n = 6 tumors per group, mean ± SD, unpaired t-test, ns=not significant). (C) H&E images of tumors from (B) showing normal chow group (left) and doxycycline chow group (right). (D) Growth curves of Pa02c and Pa20c cells cultured in 10% FBS or 10%

LPDS and varying concentrations of indicated statins for 72 h (n = 3 biological replicates, mean ± SD). (E) Tumor volumes of Pa16c subcutaneous xenografts from mice treated with vehicle (water) or 20 mg/kg Fluvastatin intraperitoneally (n = 10 mice per group, mean ± SD). (F) Tumor weights on day 29 from vehicle- or Fluvastatin-treated mice shown in (E) (n = 10 mice per group, mean ± SD, unpaired t-test, ns=not significant). Individual tumors are shown in right panel.

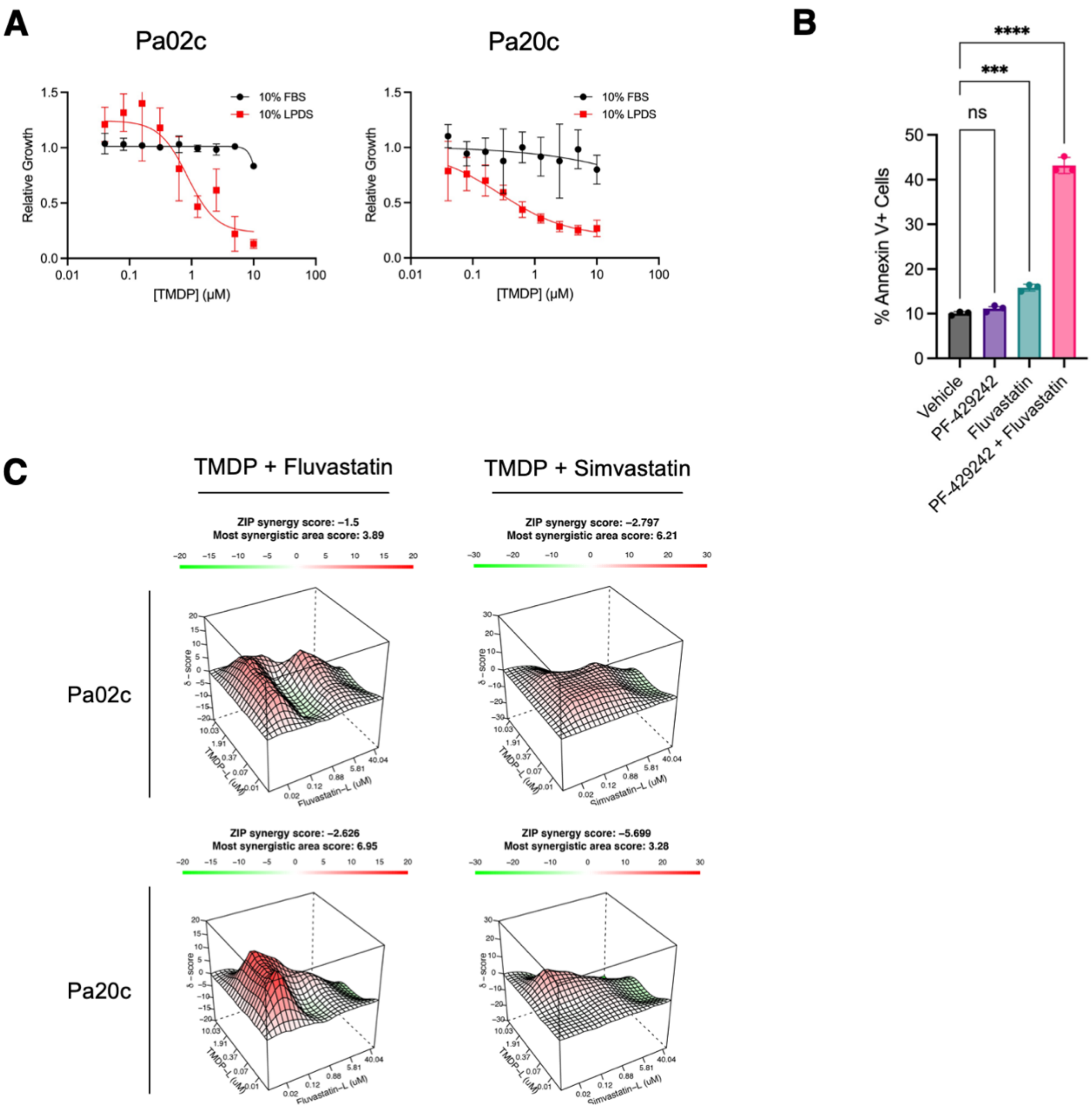


**Figure S3.** Statins induce apoptosis synergistically with SREBP inhibitors. (A) Growth curves of Pa02c and Pa20c cells cultured in 10% FBS or 10% LPDS and varying concentrations of TMDP for 72 h (n = 3 biological replicates, mean ± SD). (B) Annexin V flow cytometry quantification of Pa03c cells cultured in 0.5 μM PF-429242 and/or 5 μM Fluvastatin in 10% LPDS for 48 h (n = 3 biological replicates, mean ± SD, One-way ANOVA, ***p<0.001, ****p<0.0001). (C) 3D synergy landscape of Pa02c (top panel) and Pa20c (bottom panel) cells treated with either TMDP + Fluvastatin or TMDP + Simvastatin (n = 2 biological replicates for each dose. Each map shows the synergistic (red) and antagonistic (green) dose regions. The most synergistic area score is the average score of the highest 3x3 dose-window average across the entire matrix. A positive score indicates synergy while a negative score indicates antagonism.

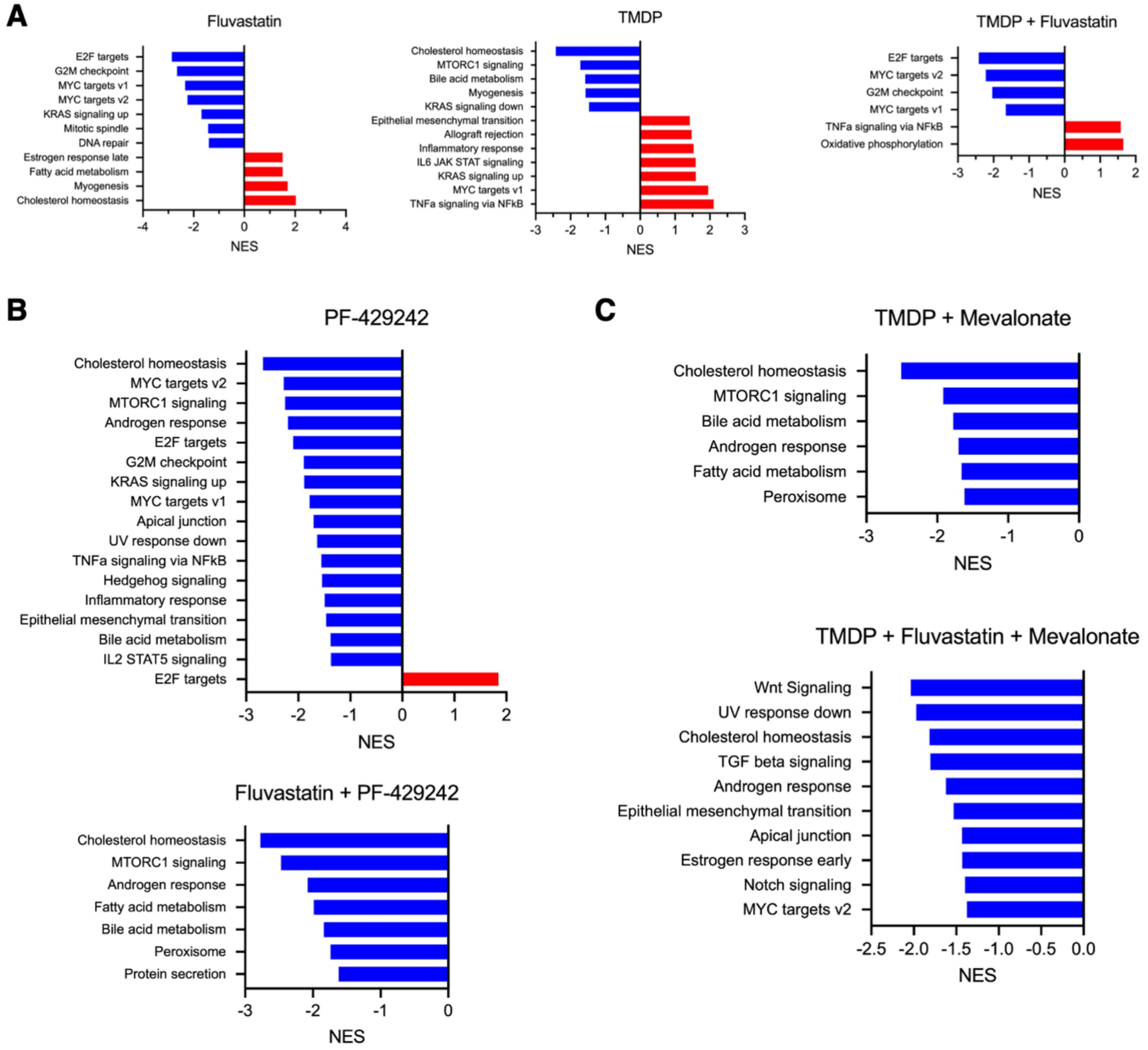


**Figure S4.** Bulk RNA-seq reveals broad cellular stress in response to combination drug treatment. (A) Identical to Fig. 4A to assist with readability. NES scores for Hallmark gene sets with FDR < 0.1 of RNA-seq samples from cells cultured in 5 μM Fluvastatin, 2.5 μM TMDP, or the drug combination in 10% LPDS for 16 h. (B) NES scores for Hallmark gene sets with FDR < 0.1 of RNA-seq samples from Pa03c cells treated with 1 μM PF-429242 and/or 5 μM Fluvastatin in 10% LPDS for 16 h. (C) NES scores for Hallmark gene sets with FDR < 0.1 of RNA-seq samples from Pa03c cells treated with 2.5 μM TMDP and/or 5 μM Fluvastatin in 10% LPDS supplemented with 1 mM mevalonate for 16 h.

### Materials and Methods

*Chemical Reagents*

We obtained chemicals from the following manufacturers: cholesterol (MilliporeSigma, C3045), LDL (Prospec Bio, PRO-562), oleic acid-albumin (MilliporeSigma O3008), mevalonate (MilliporeSigma, M4667). We obtained drug reagents from the following manufacturers: PF-429242 dihydrochloride (MedChemExpress #HY-13447A), N2,N2,N6,N6-tetrakis[2-methoxyethyl]-4,8di[piperidin-1-yl]pyrimido[5,4-d]pyrimidine-2,6-diamine (TMDP, generously provided by Dr. Tim Osborne, Johns Hopkins All Children's Hospital), Atorvastatin (MilliporeSigma #PZ0001), Fluvastatin (MilliporeSigma #SML0038), Simvastatin (MilliporeSigma #S6196), Rosuvastatin (MilliporeSigma #SML1264), Pravastatin (MilliporeSigma #P4498), GGTI-298 (Cayman #161761), FTI-277 (SelleckChem #S7465), YM-53601 (Cayman #18113), puromycin (MilliporeSigma #P8833), blasticidin (Corning #30-100-RB). For all in vitro studies, compounds were dissolved in DMSO. For in vivo studies, Fluvastatin was dissolved in sterile water.

*CRISPR Plasmid Library Generation*

A list of known SREBP target genes was compiled from published literature (1,2). Ten sgRNA sequences for each gene were chosen from published CRISPR knockout libraries (3,4) accessible at https://www.addgene.org/crispr/libraries. The plasmid library was constructed following a protocol previously published by Wang et al. (5) using lentiGuide-Puro (Addgene #52963) as the backbone vector. The custom oligo pool was synthesized by CustomArray Inc. Plasmid library representation was validated using Illumina sequencing. MAGeCK-VISPR (v.0.5.6) was used to map counts back to original plasmid library sequences.

*Mouse Husbandry*

The Johns Hopkins University animal care and use program is accredited by AAALAC International. All mouse experimental procedures were reviewed and approved by Johns Hopkins

Institutional Animal Care and Use Committee. Routine health surveillance using dirty bedding sentinel serology indicated that the mice were free of the following organisms: mouse hepatitis virus, minute virus of mice, mouse parvovirus, epizootic diarrhea of infant mice (rotavirus), Theiler's murine encephalomyelitis virus, murine norovirus, Sendai virus, pneumonia virus of mice, reovirus, lymphocytic choriomeningitis virus, ectromelia virus, mouse adenovirus (FL & K87), mouse cytomegalovirus, *Mycoplasma pulmonis*, fur mites, and pinworms. Mice were housed in social groups (2-5 mice) of the same sex in individually ventilated cages (Allentown Caging Inc., Allentown, NJ) with autoclaved corncob bedding (Teklad, Envigo, NJ) and nesting material (Animal Specialties and Provisions, Quakertown, PA). Mice are visually assessed every morning, and cages were changed every 14 days. Autoclaved feed (Teklad Global 2018S or Teklad Global 2018, 625 Doxycycline, #TD.01306 where specified) and water were provided *ab libitum*, and water was provided via an in-cage automated watering system (Systems Engineering, Inc., Napa, CA). Doxycycline diet was replaced every 3 days to maintain freshness. The room was maintained at 22±1°C on a 14:10 light:dark cycle at 40-70% humidity. Euthanasia was performed using either carbon dioxide asphyxiation at 30-70% volume displacement or isoflurane anesthesia followed by either cervical dislocation or exsanguination and bilateral thoracotomy.

*Tumor Genomic DNA Isolation*

Tumors were minced and incubated in 8 mL SDS lysis buffer (100 mM NaCl, 50 mM Tris (pH 8.1), 5 mM EDTA, 1% (w/v) SDS) per ~1 g of tumor and 0.25 mg/mL Proteinase K (Qiagen #19133) overnight at 55˚C with rocking. Lysates were centrifuged at 2,500 x g for 10 min, and 1 mL lysate was used for phenol chloroform protein extraction. Phenol:chloroform:isoamyl alcohol (1 ml, Fisher #BP1752I-400) was added, vortexed for 30 sec, and then centrifuged at 5,000 x g for 5 min. The aqueous layer was carefully removed and transferred to a new tube. These steps were repeated for a total of 3 times to remove proteins from sample. 0.1 mL 3 M sodium acetate (pH 5.0) was then added. To precipitate DNA, 0.7 mL isopropanol was added and centrifuged at

15,000 x g for 15 min at 4˚C. The pellet was washed twice with 70% ethanol and centrifuged at 15,000 x g for 5 min at 4˚C. Pellet was air dried and dissolved in ~250 µL EB buffer (Qiagen #19086) containing 100 µg/mL RNase A (Thermo #EN0531).

*Generation of CRISPRi-knockdown cells*

TLCV2 (Addgene #87360) and dCas9-KRAB-MeCP2 plasmids (Addgene #110821) were used for the plasmid backbone. Guide RNA sequences (**Table S2**) were cloned into the TLCV2 vector after Esp3I (New England Biolabs #R0734S) digestion. The dCas9-KRAB-MeCP2 cassette was amplified by PCR. The Cas9 sequence was excised from the TLCV2-sgRNA plasmid by digestion with AgeI and BamHI. Ligation of the dCas9-KRAB-MeCP2 PCR fragment with the TLCV2 backbone was done using the In-Fusion Snap Assembly Master Mix (Takara #638909) following the manufacturer's protocol. Pa03c cells were infected with dCas9 virus. After selection with 1.5 µg/mL puromycin for 72 h, polyclonal cells were treated with 1 µg/mL doxycycline for 72 h. Following treatment, tdTomato-positive and GFP-positive cells were selected using the Sony SH800 cell sorter. Cells were then dilution cloned in 96-well plates to isolate monoclonal populations and screened for dox-inducible knockdown by qPCR and western blot.

*Immunoblot Analysis*

Cells were washed twice with PBS and then lysed in ~0.1 mL (per 0.5 x $10^6$ cells) SDS lysis buffer (10 mM Tris-HCl (pH 6.8), 100 mM NaCl, 1 mM EDTA, 1 mM EGTA, 1% (w/v) SDS) supplemented with protease inhibitors (5 µg/mL pepstatin A, 10 µg/mL leupeptin, 0.5 mM PMSF, 1 mM DTT). Cell lysate was briefly probe sonicated to shear DNA and then heated at 95˚C for 5 min. For membrane protein analysis, Mem-PER Plus Membrane Protein Extraction Kit was used following the manufacturer's protocol (ThermoFisher #89842). Protein concentration was determined by BCA Protein Assay Kit (Pierce #100389) with BSA standards following the manufacturer's protocol. Proteins were separated by SDS-PAGE and transferred to nitrocellulose membranes by semi-dry transfer in the BioRad Trans-Blot Turbo Transfer system. Membranes were blocked in 5% (w/v) non-fat milk in PBST (PBS with 0.5% (v/v) Tween-20) for 30 min at room temperature. Primary antibodies were diluted in PBST and incubated at 4°C overnight. Membranes were washed 3 times in PBST at room temperature. Secondary antibodies were incubated at room temperature for 1 h followed by 3 washes in PBST at room temperature. Detection of fluorescence was performed using Odyssey CLx imaging system (LI-COR Biosciences).

*Antibody Information*

The following antibodies were used: HMGCR (1:100, Millipore #MABS1233), Calnexin (1:500, Millipore #20880), Rac1/2/3 (1:500, Santa Cruz Biotechnology #sc-514583), RhoA (1:500, Santa Cruz Biotechnology #sc-418), Rheb (1:500, Cell Signaling Technology #13879S), Kras (1:500, Sigma #WH0003845M1), E-cadherin (1:500, Cell Signaling Technology #3195T), GGPS1 (1:500, Santa Cruz Biotechnology #sc-271679), Phospho-p70 S6 Kinase (1:1000, Cell Signaling Technology #2708T), p70 S6 Kinase (1:1000, Cell Signaling Technology #2708T), Phospho-p44/42 MAPK (1:1000, Cell Signaling Technology #4370T), p44/42 MAPK (1:1000, Cell Signaling Technology #4695T), Phospho-p38 (1:1000, Cell Signaling Technology #4511T), p38a MAPK (1:1000, Cell Signaling Technology 9228S), GAPDH (1:5000, Sigma #G9545), Actin (1:500, Santa Cruz Biotechnology #SC-47778), IRDye 680RD goat anti-mouse (1:20,000, Fisher #NC0252290) and anti-rabbit (1:20,000, Fisher #NC0252291), IRDye 800CW goat anti-mouse (1:20,000, Fisher #NC9401841) and anti-rabbit IgG (1:20,000, Fisher #NC9401842).

*Bioinformatic Analyses*

For CRISPR knockout screening analysis, MAGeCK-VISPR (v.0.5.6) was used for read mapping, quality control, and analysis as described in the MAGeCK manual (https://sourceforge.net/p/mageck/wiki/Home/). MAGeCK MLE was used for multiple sample comparisons of FBS and LPDS samples and separate analyses were conducted for each time point. MAGeCK RRA was used for in vivo samples. In both cases, day 0 library-infected cell samples were used as control reference. MAGeCKFlute (v. 1.10.0) was used for visualization of the screening results.

For RNA-seq, reads were trimmed using Trimmomatic (v.0.36) and then mapped to the *Homo sapiens* reference genome GRCh38 using STAR (v.2.5.2b). Exonic reads were counted using FeatureCounts (v.1.5.2) and EdgeR (v.3.32.1; R v.4.1.3) was used for differential gene expression analysis. Gene set enrichment analysis (GSEA) was conducted using MSigDB (v.4.1.0) (6). An ordered log(fold change) ranked list was provided as input to determine Hallmark gene set enrichment (7).

TNMplot was used to compare pancreas expression of *GGPS1* from normal and tumor (www.TNMplot.com) (8). Data were downloaded and plotted.
